# DxFit: An ensemble method for identifying EHR diagnoses consistent with a molecular finding

**DOI:** 10.64898/2026.04.24.720629

**Authors:** Rebecca I. Torene, Karyn Meltz Murphy, Tracy Brandt, Kyle Retterer

## Abstract

As population DNA sequencing becomes more common, genomic-first approaches are increasingly used to identify individuals with possible rare genetic disorders. To accurately estimate prevalence and penetrance, these studies often confirm manifestation of the disorder using electronic health records (EHRs). Multiple strategies exist to search the EHR for diagnoses of rare disorders, however, each has its limitations. We have developed a portable, ensemble tool, DxFit, that mines EHR data (ICD codes and structured diagnosis descriptions from billing code and problem list tables) for a diagnosis consistent with a given rare genetic disorder. DxFit combines evidence across four strategies: (1) gene name searches in diagnosis descriptions and notes, (2) ICD conversion to Mondo rare disorder ontology to find exact and nearby matches, (3) word embedding similarity searches, and (4) Jaccard similarity matches. DxFit prioritizes the match type and outputs the most confident match for each participant-disorder pair. On a cohort of 350 participants with a known positive result from diagnostic genetic testing for developmental disorders, DxFit had a sensitivity of 88.7% and specificity of 86.2% using default parameters. Adjusting the linguistic scoring thresholds from 0.8 to 0.7 and allowing for synonymous matches yielded a sensitivity of 92.7% and specificity of 84.5%. Partitioning EHR evidence into windows before and after genetic testing demonstrates, as expected, that the overall DxFit rates increase after testing and the match types become more confident. DxFit is available to the public and has extensive customization options to support a wide range of uses.

**Graphical Abstract:** 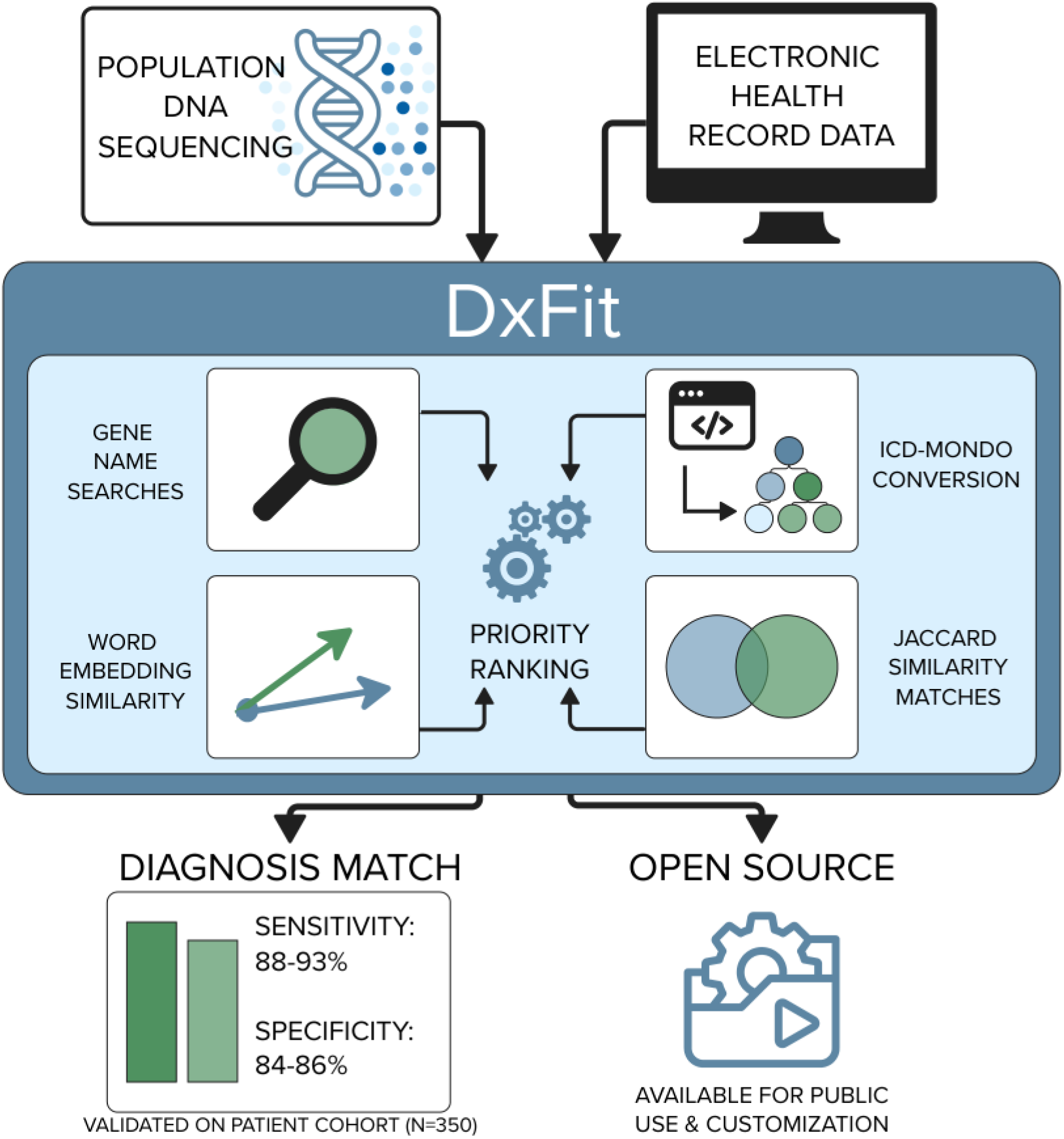

## Main

Our collective understanding of rare genetic disorders comes largely from clinically ascertained cohorts, following a phenotype-first approach in which individuals with relevant features are referred for genetic testing^1^. Recently, as large, population-based cohorts have undergone DNA sequencing, genomic-first studies have mined genomic data for pathogenic variants regardless of whether the participant exhibits relevant clinical features^2–5^. To accurately estimate variant penetrance and evaluate specificity of these population-based studies, it is important to determine whether the participant has evidence suggestive of the genetic disorder. This evidence is often sought via electronic health record (EHR) data and can be a specific diagnosis or a collection of phenotypes consistent with a genetic disorder.

Previously, we published a genomic-first approach to identify high confidence pathogenic and likely pathogenic variants that cause known genetic conditions in a healthcare-based cohort of 216,000 participants^6^. In doing so, we identified a need for a method to match an individual’s suspected rare genetic disorder to the clinical diagnoses in their EHR. Prior studies have addressed this need by creating EHR-to-Mondo maps^7,8^, using semantic similarity tools to match disorder names to diagnosis descriptions^9^, evaluating confirmatory laboratory results for metabolic disorders^3^, or by manual review^10^. These approaches all have their strengths and limitations.

Direct mapping of EHR-based controlled vocabularies to Mondo can suffer from both sparsity (under-matching) and greedy matching (over-matching) depending on the mapping strategy. The Monarch Institute, which developed the Mondo ontology, provides direct maps from International Classification of Diseases (ICD) to Mondo; however, this map is sparse with 75.1% of Mondo terms having no ICD mapped. Making use of the ontology, 32.6% of Mondo terms have an ICD mapped within two degrees, leaving 67.4% unmapped. This is not unexpected, as these two vocabularies have different purposes and levels of specificity with ICD being used for medical billing and Mondo being used to delineate rare genetic disorders.

Conversely, EHR-to-Mondo maps can suffer from greedy matching where a single ICD code maps to multiple Mondo IDs. The *omop2obo*^7^ approach maps SNOMED terms to Mondo without this issue. However, converting ICD to SNOMED to use *omop2obo* introduces one-to-many mappings: a single ICD code can map to multiple SNOMED codes which then map to multiple Mondo disorders. On average, each ICD code will map to 8.1 SNOMED terms. For example, a single ICD code R56.9 for convulsive seizures maps to 108 SNOMED terms which, in turn, map to 77 Mondo IDs. This is not unexpected as this term is a phenotype and is not specific to a single rare genetic disorder. ICDs that are more specific will map to fewer SNOMED terms, such as G71.01 for Duchenne’s or Becker muscular dystrophy which maps to 12 Mondo IDs, all of which are closely related myopathy disorders.

Prior studies have attempted to overcome these mapping issues by using semantic similarity instead^9^. Semantic similarity can be used to both prioritize one-to-many mappings described above and to identify new mappings between vocabularies. Previously, we observed a high rate of semantic similarity matches among random controls^6^. Upon review, we observed false-positive matches caused by nonspecific disease descriptors (e.g., “abnormal” or “type”) and by rare genetic disorder matches to their common counterparts (e.g., “Maturity-Onset Diabetes of the Young” matching to “type II diabetes” or “inherited obesity” matching to “obesity”). Likewise, we also observed that eponymous and acronym disorders were missing from standard natural language processing (NLP) models and thus would not produce semantic similarity scores unless there was an exact string match.

To address the lack of a method for detecting rare genetic disorders in a patient’s EHR that is agnostic to gene and disorder type (e.g., metabolic, skeletomuscular, autoimmune), we created an ensemble method, DxFit. DxFit uses a patient’s diagnoses, in the form of ICD codes, brief descriptions, and patient notes, and searches for specific pieces of evidence of a diagnosed or suspected rare genetic disorder. Since we originally published this method, we have made the tool available to the public, added the ability to use clinical notes, generalized the disease vocabulary that can be used, and performed benchmarking on a positive control dataset.

DxFit assesses evidence of a rare disorder in the EHR using four approaches (Figure 1, Table 1):

**Table 1.**
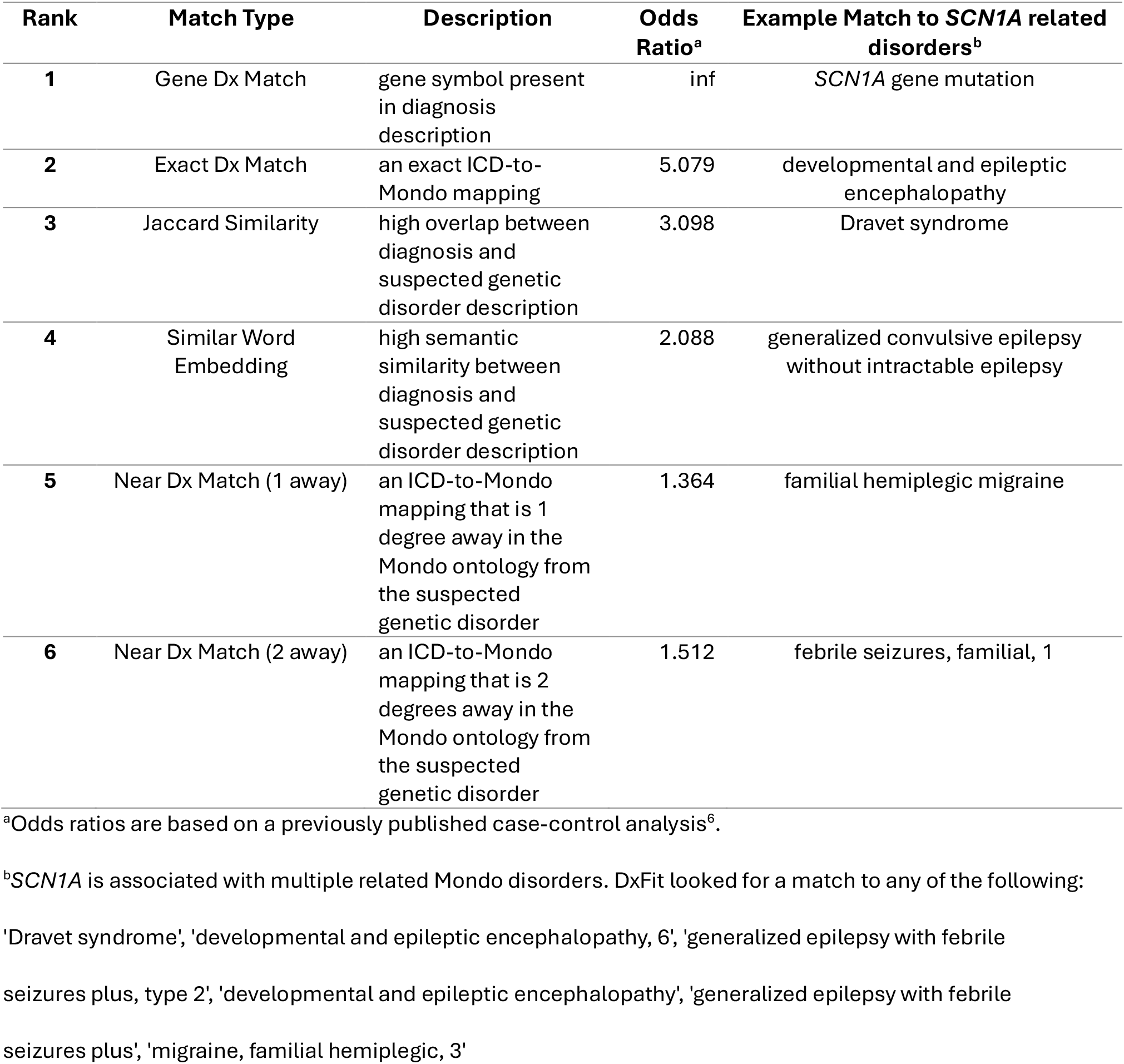
DxFit match types.

| Rank | Match Type | Description | Odds Ratio<sup>a</sup> | Example Match to *SCN1A* related disorders<sup>b</sup> |
| --- | --- | --- | --- | --- |
| 1 | Gene Dx Match | gene symbol present in diagnosis description | inf | *SCN1A* gene mutation |
| 2 | Exact Dx Match | an exact ICD-to-Mondo mapping | 5.079 | developmental and epileptic encephalopathy |
| 3 | Jaccard Similarity | high overlap between diagnosis and suspected genetic disorder description | 3.098 | Dravet syndrome |
| 4 | Similar Word Embedding | high semantic similarity between diagnosis and suspected genetic disorder description | 2.088 | generalized convulsive epilepsy without intractable epilepsy |
| 5 | Near Dx Match (1 away) | an ICD-to-Mondo mapping that is 1 degree away in the Mondo ontology from the suspected genetic disorder | 1.364 | familial hemiplegic migraine |
| 6 | Near Dx Match (2 away) | an ICD-to-Mondo mapping that is 2 degrees away in the Mondo ontology from the suspected genetic disorder | 1.512 | febrile seizures, familial, 1 |

<sup>a</sup>Odds ratios are based on a previously published case-control analysis<sup>6</sup>.
<sup>b</sup>*SCN1A* is associated with multiple related Mondo disorders. DxFit looked for a match to any of the following:
'Dravet syndrome', 'developmental and epileptic encephalopathy, 6', 'generalized epilepsy with febrile seizures plus, type 2', 'developmental and epileptic encephalopathy', 'generalized epilepsy with febrile seizures plus', 'migraine, familial hemiplegic, 3'

**Figure 1.**
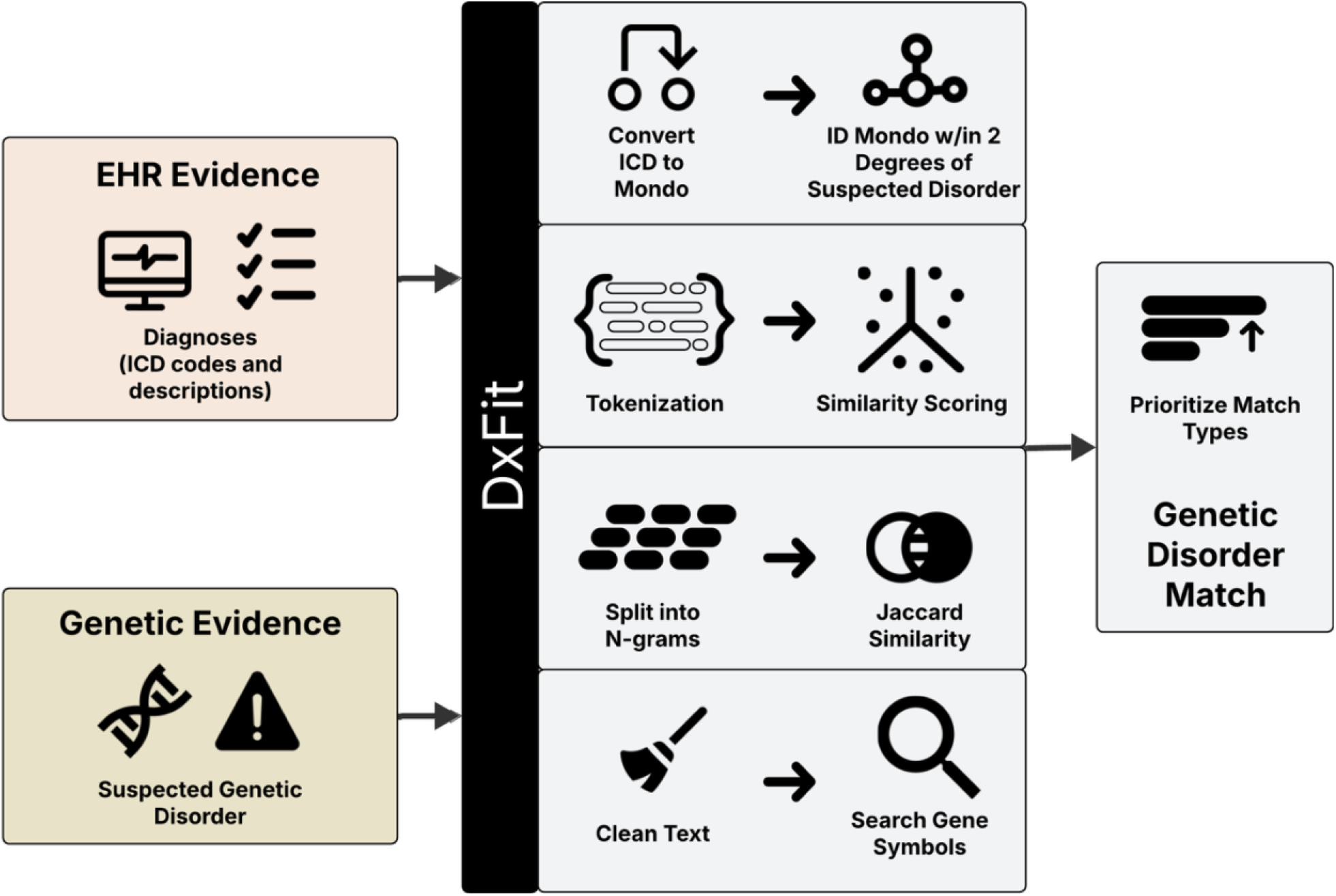
Schematic of DxFit DxFit allows a user to input a file with diagnosis codes and descriptions, medical notes (optional), and suspected rare genetic disorders. It then runs several different matching strategies to identify evidence in the EHR that is supportive of the genetic disorder. It performs *post hoc* prioritization based on prior evidence of specificity of the match type and outputs the most confident match.

1. Gene symbol search: DxFit searches the diagnosis descriptions (and notes if provided) for an exact match to the suspected disorder gene symbol
2. ICD-to-Mondo mapping: DxFit uses ICD-to-Mondo maps provided by the Monarch Initiative to identify ICDs in an individual mapping within 2 degrees of the Mondo ontology to their suspected genetic disorder
3. Similar word embedding: DxFit performs pairwise semantic similarity searches using NLP word embeddings between the suspected genetic disorder and each given diagnosis description (e.g., ICD diagnosis name typically from structured billing code and problem list EHR tables)
4. Jaccard similarity: DxFit overlaps n-grams of the suspected genetic disorder with each given diagnosis description to identify matches with high overlap

DxFit prioritizes the match type based on specificity compared to a control cohort ^6^ (Table 1).

### Problems addressed by DxFit

#### 1. ICD-to-Mondo maps having sparse or greedy matching

As described above, EHR-to-Mondo maps can be imprecise and thus lead to both under-and over-matching. DxFit minimizes this issue by starting from a sparser map (ICD-to-Mondo from the Monarch Initiative) but allowing for nearby matches in the more granular Mondo ontology. If an ICD code maps directly to a Mondo ID, DxFit will return an exact match. If, instead, an individual has an ICD that maps within 1 or 2 degrees of the suspected genetic disorder, DxFit will return a “Near Dx Match (1 away)” or “Near Dx Match (2 away)” match, respectively.

We note, however, that the Mondo ontology is not uniformly specific at different depths, and allowing for 2-degree matches could lead to overly greedy matches. The node “hereditary disease” (MONDO:0003847) has 2,014 diverse genetic disorders as direct child terms. These child terms are then siblings to each other within 2 degrees of the ontology even if they are vastly different. For example, “spinal muscular atrophy with intellectual disability” (MONDO:0010054) is only 2 degrees away from “uncombable hair syndrome 1” (MONDO:0020736), an unrelated disorder. To prevent over-matching of ICD-to-Mondo disorders within 2 degrees, we custom prune the Mondo ontology used within DxFit to:

- Keep only descendants of the root term for human disease (MONDO:0700096) and edge type of “is_a”
- Remove a curated list of vague disorders including “hereditary disease”
- Remove edges within 2 degrees of the root term for human disease as these are likely system level terms

These efforts help prevent distant and overly ambiguous matches to ICD codes.

#### 2. Over-matching of semantic similarity

In our previous study, we noted an excess of controls with semantic similarity matches. To address these spurious matches, we employed three approaches: 1) omit nonspecific terms from similarity scoring, 2) omit diagnoses that are not present in the participant, and 3) drop similarity matches *post hoc* that include common disorder terms. To omit nonspecific terms, we created a custom list of stop-words to be added to the NLP model that would be ignored by the semantic similarity search algorithm. The custom stop-words are those commonly seen in diagnosis descriptions and could lead to spurious matches such as “familial”, “abnormal”, or “type”. To omit diagnoses that may not be present in the individual, we discard all diagnosis descriptions that contain any of the terms “gestational”, “family history”, or “screening”. We also observed that spurious matches among controls were due to rare forms of a genetic disorder matching to their common counterpart (e.g., periodic fever syndromes matching to acute fever). For such cases, we created a custom list of terms for which linguistic matches (either semantic or Jaccard similarity) would be omitted *post hoc*, but gene name and ICD-to-Mondo mapping matches would still be permitted. The three lists of terms used to increase specificity for DxFit (custom stop-words, discard diagnoses terms, and common linguistic matching terms) are all pre-populated in DxFit with the option for the user to modify as needed.

#### 3. Lack of eponymous and acronym disorder matching

Depending on the NLP model being used, whether it has been trained on a medical corpus, and the number of tokens in the model, it is possible that some rare genetic disorders are not adequately represented in the vector. When a disorder is missing from an NLP vector or only sparsely represented in the training data, semantic similarity tools will have difficulty finding a match. For the model we used (`en_core_sci_md`^11^), we observed that disorders missing from the NLP model vector were more likely to have eponymous or acronym-derived names. To identify matches to these disorders, we added Jaccard similarity scoring to DxFit. The suspected rare genetic disorder names and the diagnosis descriptions are split into tiled n-grams and then assessed for overlap. This allows for matching of disorder names that are otherwise missing or under-represented in the NLP model.

### Evaluating DxFit on a positive control cohort

Previously, we applied DxFit on a cohort of individuals with genomic-first findings across 490 known genetic disorders^6^. We observed a DxFit rate of 15.0%. It is unclear, however, how many of those without a DxFit match are due to lack of a confirmed diagnosis in the EHR (DxFit true negative) or if a confirmed diagnosis exists and DxFit is unable to detect it (DxFit false negative). Likewise, since DxFit allows for both exact and fuzzy matches, the observed specificity was unclear. To assess DxFit’s sensitivity and specificity, we utilized a retrospective cohort of individuals with suspected genetic developmental disorders who were referred for clinical genetic testing. Data from individuals with molecular positive findings were collected from consented participants in an IRB-approved study protocol. Control data were collected from a deidentified research database. Of the 3,380 individuals with genetic testing, 362 had a monogenic molecular diagnosis reported. We further filtered to individuals with at least one ICD code in the past 5 years, indicating relatively recent care in our health system (N=350 individuals). This development disorder cohort had a median age of 12.1 (IQR 9.4-16.3) years, was 37.1% female, and 79.4% non-Hispanic white. The median age at genetic testing was 5.8 (IQR 3.9-9.7) years. A few individuals had more than one molecular diagnosis leading to N=355 individual-disorder pairs. We also created an age and sex-matched control cohort with age matching within 2 years and 10 controls per case. Each control individual was assigned the same molecular finding as their matched case to determine whether spurious positive matches to their EHR would occur.

We evaluated all 355 single-gene, molecular positive individual-disorder pairs and their matched controls using DxFit. We used a curated set of genetic disorders from the GenCC coalition^12^ with strong or definitive evidence to map gene findings to Mondo disorders. We then processed these data through DxFit, comparing ICD codes (from diagnoses and problem lists) and their descriptions to the Mondo disorders. To represent the most likely usage of DxFit, we did not include clinical notes in this evaluation. Of the 355 individual-disorder pairs, 315 (88.7%) had at least one DxFit match (Figure 2) and the specificity was 86.2%. The majority (475/484, 98.1%), of the DxFit matches in controls (false positives) were Near Dx Match (1 or 2 away). These matches are the least confident that DxFit returns and are prioritized below gene name match, exact ICD match, and linguistic matches. The most frequent match type among cases was gene name matching to ICD descriptions (N=170, 47.9% of the total cohort). Since gene name matching is the simplest match type and does not require the technical overhead of DxFit’s ensemble methods, we evaluated how well DxFit would perform if we ignore this match type. If we omit gene name matching, we still observe that 280 individuals (79.2%) have at least one other DxFit match type (Figure 2). Similarly, 145 (40.8%) of the cohort had DxFit matches that would not have been found by gene name searching alone.

**Figure 2.**
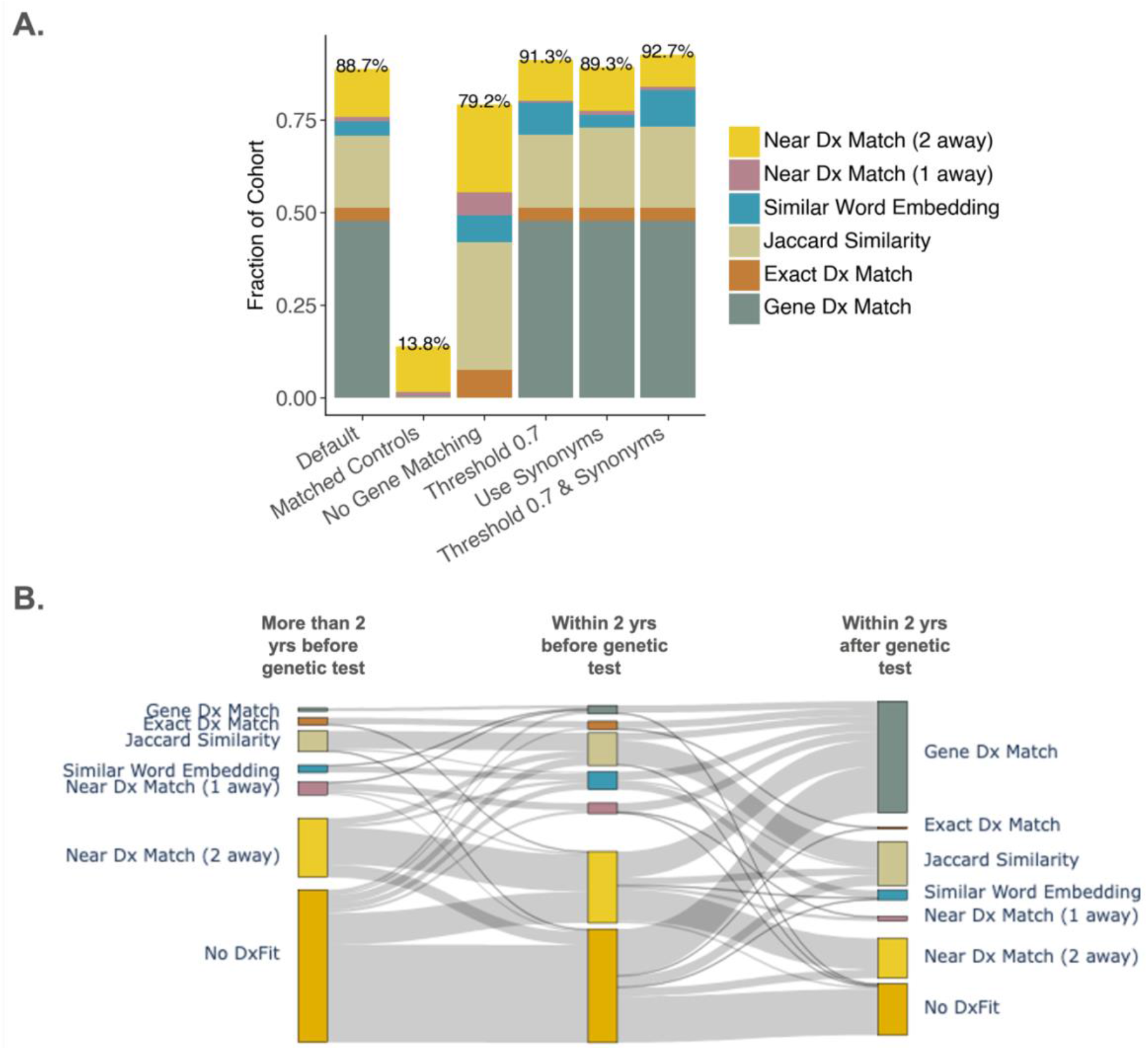
DxFit match rate on a positive control cohort A) DxFit had a sensitivity of 88.7% on a cohort of individuals with confirmed, single-gene, molecular diagnoses (N=355). Changing parameters for DxFit affects the total match rate. B) Different rates of DxFit categories across time for individuals with at least 1 diagnosis code in each of the 3 time windows (N=250). Note how the “No DxFit” category shrinks over time while confident match types like “Gene Dx Match” grow.

Of the 40 individuals with no DxFit, 9 had semantic similarity scores ≥0.7. By default, DxFit uses a conservative similarity score threshold of 0.8 for both Jaccard and word embedding semantic similarity; however, for some use cases it would be reasonable to lower this threshold to 0.7. For example, pathogenic variants in the *SETD1B* gene are associated with MONDO:0033559 (intellectual developmental disorder with seizures and language delay) which has a similarity score of 0.75 to the term “intellectual disability”. If we lower both linguistic matching thresholds from 0.8 to 0.7, the DxFit rate increases 2.6% from 88.7% to 91.3% while the specificity drops to a lesser extent (85.6%) (Figure 2).

For the remaining 31 individuals without a DxFit, we conducted manual EHR reviews. The gene name was always present in the notes, and the disorder name was also almost always present; however, sometimes the disorder name did not match the Mondo name. For example, *KDM5B* is associated with MONDO:0020850 (“intellectual disability, autosomal recessive 65”). A patient with a variant in *KDM5B* had developmental delay and learning disability listed in diagnoses but not the words, “intellectual disability” or the gene name leading to no ICD matches and low linguistic matching scores. This highlights the limitations of using ICD codes and descriptions to identify rare genomic disorders in the EHR as evidence of a rare disorder might be present in notes and other EHR fields.

To account for likely matches that might not meet linguistic similarity thresholds, DxFit has the option to include Mondo disorder synonyms. When we enable this feature, two additional positives are recalled and the DxFit rate goes from 88.7% under default parameters to 89.3% while the specificity is 85.9%. If we enable both Mondo synonym matching and lower the linguistic matching thresholds to 0.7, then the DxFit rate goes up to 92.7% in cases and 15.5% in controls, a specificity of 84.5%.

We also performed expert review of the non-exact DxFit findings in the molecular positive cases to determine poor matches with two independent reviewers. Of the 315 cases with a positive DxFit finding, 19 (6.0%) were considered poor matches to the EHR by the reviewers and 42 (13.3%) were found to match vague diagnoses in the EHR that may not be indicative of a specific genetic disorder (e.g., “intellectual disability”). All 19 of the poor matches were “Near Dx Match (2 away)” which is the least confident category DxFit reports and is prioritized last. Users of DxFit can decide to ignore this category if they would like more conservative results. The vague matches were a mix of DxFit types (Near Dx Match (2 away) – 24, Near Dx Match (1 away) – 2, Jaccard Similarity – 10, Similar Word Embedding – 6). Although vague, these EHR findings are consistent with the suspected genetic disorder.

Most molecular-positive participants had a documented genetic testing date in the EHR (N=341 participants) and 334 participants had at least 1 ICD code before and after their testing date. By splitting participant ICDs into before and after genetic testing dates, we see that both the rate of DxFit and the level of confidence of DxFit increase significantly after a confirmed molecular diagnosis (DxFit rate from 56.8% to 80.4%, Fisher p=1.73e-08, Figure 2). We also observe that rates of DxFit increase in the 2 years prior to genetic testing as clinical suspicion of a genetic disorder leads to genetic. Using, the prioritization of DxFit match types by confidence level, we evaluated the change in diagnostic confidence before versus after genetic testing. Of 334 participant-disorder pairs where the participant had at least one ICD code before and after genetic diagnosis, 182 (54.5%) participant-diagnosis pairs increase in confidence by DxFit type after genetic testing. For example, 5 participants with a molecular diagnosis of *CACNA1A*-related complex neurological disorder (MONDO:0100254) had a DxFit result of Near Dx Match (2 away) due to diagnosis code ICD9 315.39 (“Other developmental speech or language disorder”). Individuals with *CACNA1A* pathogenic variants often experience speech and language delays which likely led to this ICD diagnosis code being added to their EHR shortly prior to genetic testing. In the two years after genetic testing, 3/5 of these individuals had a more specific genetic diagnosis added to their EHR that included the gene name in the diagnosis description. DxFit detects both the vaguer diagnoses that support the presence of a rare genetic disorder prior to confirmed molecular diagnosis and the more confident diagnosis after genetic testing.

### Customization Options

To make DxFit useful to the broader genomics community, we have added numerous customization options:

- **NLP model** – DxFit has thus far only been evaluated with the `en_core_sci_md` model, however, other NLP models including those with larger corpora and specialized training on medical text may be substituted.
- **Evaluate one, or more disorders per individual** – The input data format does not assume a one-to-one mapping of individual to suspected genetic disorder. Instead, the user may evaluate a collection of disorders together or as separate rows in the input file to evaluate each disorder individually. This is useful when multiple related Mondo disorders represent a phenotype spectrum of a single disorder versus when a participant has multiple significant genetic findings that could manifest in distinct disorders.
- **Input disorder names/descriptions** – Thus far, we have compared ICD-to-Mondo disorders, however, the user can use their own rare disorder vocabulary such as OMIM or Orphanet names instead. In such cases, gene name searching, semantic and Jaccard similarity will still be evaluated, but no mapping from ICD will occur to other vocabularies. Similarly, if disorder names are not provided, DxFit will look them up by Mondo ID. There is an additional flag the user can provide to match to Mondo synonyms as well.
- **Semantic and Jaccard scoring thresholds** – For our prior efforts in evaluating across a large cohort and diverse genetic disorders, we used a stringent score threshold of 0.8 for both semantic and Jaccard similarity. For more targeted efforts, or when using DxFit as a precursor to manual review, it would be reasonable to lower these thresholds for one or both match types.
- **Control cohorts** – We allow the user to submit a control cohort by having a “cohort” label in the disorder file. This will compare DxFit rates by cohort label in the stacked bar plot that DxFit produces. Alternatively, the user can apply the `create_control_cohort` flag which will shuffle the individual IDs assigned to genetic disorders as a negative control cohort.
- **Update ICD-to-Mondo maps** – For the user’s convenience, we provide a map of ICD codes to Mondo IDs which was obtained from the Monarch Initiative’s API. If the user would like to update this map, we provide an option to do so automatically.
- **Modify stop-words, terms omitted from linguistic matching, and common disorder terms** – To improve the specificity of semantic and Jaccard similarity, there are three lists of terms used to clean the data at various steps in the matching process. Depending on the disease area being evaluated, the user can modify these terms to allow for more sensitive or specific matches.

### Summary

DxFit is an ensemble tool used to match rare genetic disorders to diagnoses in the EHR at scale. It is agnostic to medical specialty and has 88.7-92.7% recall on known positively diagnosed individuals. It can be adapted to use different NLP models and vocabularies to suit user needs. It is our hope that those performing genomic-first studies, can use this tool to aid in their research and prioritize manual review.

## Acknowledgments

The authors would like to thank Melissa A. Kelly and Hunt Willard for helpful discussions during DxFit development. We thank the patients and their families who enrolled in this study from the Department of Developmental Medicine for their participation and time.

## Data and code availability

DxFit is available at https://github.com/GeisingerResearchPublic/DxFit

## eTOC blurb

DxFit is an ensemble tool that mines electronic health record data to identify evidence of rare genetic disorders. Combining multiple strategies, DxFit achieved up to 92.7% sensitivity in a positive control cohort and offers extensive customization for diverse genomic-first studies.

## References

1. Bamshad, M.J., Nickerson, D.A., and Chong, J.X. (2019). Mendelian Gene Discovery: Fast and Furious with No End in Sight. The American Journal of Human Genetics 105, 448–455. 10.1016/J.AJHG.2019.07.011.

2. Bates, B.A., Bates, K.E., Boris, S.A., Wessman, C., Stone, D., Bryan, J., Davis, M.F., and Bailey, M.H. (2025). Intersection of rare pathogenic variants from TCGA in the All of Us Research Program v6. Human Genetics and Genomics Advances 6. 10.1016/j.xhgg.2025.100405.

3. Goodrich, J.K., Singer-Berk, M., Son, R., Sveden, A., Wood, J., England, E., Cole, J.B., Weisburd, B., Watts, N., Caulkins, L., et al. (2021). Determinants of penetrance and variable expressivity in monogenic metabolic conditions across 77,184 exomes. Nat. Commun. 12. 10.1038/S41467-021-23556-4.

4. Blair, D.R., and Risch, N. (2024). Dissecting the Reduced Penetrance of Putative Loss-of-Function Variants in Population-Scale Biobanks. medRxiv, 2024.09.23.24314008. 10.1101/2024.09.23.24314008.

5. Forrest, I.S., Chaudhary, K., Vy, H.M.T., Petrazzini, B.O., Bafna, S., Jordan, D.M., Rocheleau, G., Loos, R.J.F., Nadkarni, G.N., Cho, J.H., et al. (2022). Population-Based Penetrance of Deleterious Clinical Variants. JAMA 327, 350–359. 10.1001/JAMA.2021.23686.

6. Torene, R.I., Murphy, K.M., Brandt, T., Kelly, M.A., Willard, H.F., and Retterer, K. (2025). A scalable approach for genomic-first rare disorder detection in a healthcare-based population. The American Journal of Human Genetics 112, 2565–2577. 10.1016/j.ajhg.2025.09.010.

7. Callahan, T.J., Stefanski, A.L., Wyrwa, J.M., Zeng, C., Ostropolets, A., Banda, J.M., Baumgartner, W.A., Boyce, R.D., Casiraghi, E., Coleman, B.D., et al. (2023). Ontologizing health systems data at scale: making translational discovery a reality. 10.1038/s41746-023-00830-x.

8. Shefchek, K.A., Harris, N.L., Gargano, M., Matentzoglu, N., Unni, D., Brush, M., Keith, D., Conlin, T., Vasilevsky, N., Zhang, X.A., et al. (2020). The Monarch Initiative in 2019: An integrative data and analytic platform connecting phenotypes to genotypes across species. Nucleic Acids Res. 48, D704–D715. 10.1093/NAR/GKZ997,.

9. Herr, K., Lu, P., Diamreyan, K., Xu, H., Mendonca, E., Weaver, K.N., and Chen, J. (2024). Estimating prevalence of rare genetic disease diagnoses using electronic health records in a children’s hospital. Human Genetics and Genomics Advances, 100341. 10.1016/j.xhgg.2024.100341.

10. Gold, J.I., Kripke, C.M., Mansfield, A.J., Li, A., Lopez, A., Hawes, A., Averitt, A., Damask, A., Deubler, A., Ziyatdinov, A., et al. (2025). Exclusion-based exome sequencing in critically ill adults 18–40 years old has a 24% diagnostic rate and finds racial disparities in access to genetic testing. Am. J. Hum. Genet. 112, 1792–1804. 10.1016/j.ajhg.2025.06.007.

11. Neumann, M., King, D., Beltagy, I., and Ammar, W. (2019). ScispaCy: Fast and robust models for biomedical natural language processing. BioNLP 2019 - SIGBioMed Workshop on Biomedical Natural Language Processing, Proceedings of the 18th BioNLP Workshop and Shared Task, 319–327. 10.18653/V1/W19-5034.

12. DiStefano, M.T., Goehringer, S., Babb, L., Alkuraya, F.S., Amberger, J., Amin, M., Austin-Tse, C., Balzotti, M., Berg, J.S., Birney, E., et al. (2022). The Gene Curation Coalition: A global effort to harmonize gene–disease evidence resources. Genetics in Medicine 24, 1732–1742. 10.1016/j.gim.2022.04.017.

